# Global evolutionary dynamics and resistome analysis of *Clostridioides difficile* ribotype 017

**DOI:** 10.1101/2021.07.04.451084

**Authors:** Korakrit Imwattana, Papanin Putsathit, Deirdre A Collins, Teera Leepattarakit, Pattarachai Kiratisin, Thomas V Riley, Daniel R Knight

## Abstract

*Clostridioides difficile* PCR ribotype (RT) 017 ranks among the most successful strains of *C. difficile* in the world. In the past three decades, it has caused outbreaks on four continents, more than other “epidemic” strains, however, our understanding of the genomic epidemiology underpinning the spread of *C. difficile* RT 017 is limited. Here, we performed high-resolution phylogenomic and Bayesian evolutionary analyses on an updated and more representative dataset of 282 non-clonal *C. difficile* RT 017 isolates collected worldwide between 1981 and 2019. These analyses place an estimated time of global dissemination between 1953 and 1983 and identified the acquisition of the *ermB*-positive transposon *Tn6194* as a key factor behind global emergence. This coincided with the introduction of clindamycin, a key inciter of *C. difficile* infection, into clinical practice in the 1960s. Based on the genomic data alone, the origin of *C. difficile* RT 017 could not be determined, however, geographical data and records of population movement suggest that *C. difficile* RT 017 had been moving between Asia and Europe since the Middle Ages and was later transported to North America around 1860 (95% CI: 1622 – 1954). A focused epidemiological study of 45 clinical *C. difficile* RT 017 genomes from a cluster in a tertiary hospital in Thailand revealed that the population consisted of two groups of multidrug-resistant (MDR) *C. difficile* RT 017 and a group of early, non-MDR *C. difficile* RT 017. The significant genomic diversity within each MDR group suggests that although they were all isolated from hospitalised patients, there was likely a reservoir of *C. difficile* RT 017 in the community that contributed to the spread of this pathogen.

**Impact statement:** This study utilises genomic sequence data from 282 non-clonal *C. difficile* ribotype (RT) 017 isolates collected from around the world to delineate the origin and spread of this epidemic lineage, as well as explore possible factors that have driven its success. It also reports a focused epidemiological investigation of a cluster of *C. difficile* RT 017 in a tertiary hospital in Thailand to identify possible sources of transmission in this specific setting.

**Data summary:** All new WGS data generated in this study has been submitted to the European Nucleotide Archive under the BioProject PRJEB44406 (sample accession ERS6268756 – ERS6268798). The complete genome of *C. difficile* MAR286 was submitted to GenBank under BioProject PRJNA679085 (accession CP072118). Details of all genomes included in the final analyses are available in the **Supplementary Document**, available at 10.6084/m9.figshare.14544792.

## Introduction

*Clostridioides difficile* PCR ribotype (RT) 017, or sequence type (ST) 37, ranks among the most successful strains of *C. difficile*. Despite producing only one functional toxin (toxin B), *C. difficile* RT 017 has spread widely and caused outbreaks globally (1). The severity of *C. difficile* infection (CDI) caused by RT 017 has been comparable to infection caused by *C. difficile* strains producing two or three toxins (2–4). One factor that may have contributed to the success of *C. difficile* RT 017 is antimicrobial resistance (AMR) (5).

The evolutionary origins of *C. difficile* RT 017 remain contentious (1). Possible contributing factors included the early erroneous dismissal of *C. difficile* RT 017 as non-pathogenic due to its lack of toxin A (6), and the use of diagnostic tests that only detected toxin A (7). By the time that the pathogenicity of *C. difficile* RT 017 was recognised (1995) (8), the strain had already spread across the globe (1). Based on the geographical distribution of *C. difficile* clades, *C. difficile* RT 017, a member of evolutionary clade 4, has been hypothesised to have originated in Asia (1). This is supported by various epidemiologic studies reporting a high prevalence of *C. difficile* RT 017 and closely related clade 4 strains in Asia, especially Southeast Asia (9–13). However, most of these studies only included a few historical *C. difficile* RT 017 strains available from the region to verify this hypothesis (9–13). A 2017 study by Cairns *et al*. analysed whole-genome sequence (WGS) data from 277 *C. difficile* RT 017 strains and suggested an alternative hypothesis, that *C. difficile* RT 017 originated in North America, spread to Europe in the early 1990s and later to other regions (14). Despite the large dataset, this conclusion might have been influenced by a strain selection bias, as the North American strains included in the study were relatively older than strains from other regions. A recent study based mainly on the same global dataset agreed that *C. difficile* RT 017 spread first from North America but suggested that the spread may have happened before 1970 (15). In our study, we included a larger number of strains, with a few early European strains and a greater diversity of Asian strains. We aimed to further explore the origin and spread of *C. difficile* RT 017, as well as the key genetic factors driving its success.

## Methods

### *C. difficile* RT 017 genomes

This study included 933 *C. difficile* RT 017 strains from three collections; a set of 45 clinical *C. difficile* RT 017 strains from Thailand (32 phenotypically MDR and 13 non-MDR) some of which have been described previously (16), 102 previously unpublished *C. difficile* RT 017 strains from our laboratory’s collection and 786 *C. difficile* RT 017 genomes publicly available at the NCBI Sequence Read Archive (https://www.ncbi.nlm.nih.gov/sra/) as of January 2020. These collections included genomes of three *C. difficile* RT 017 isolated in the early 1980s (courtesy of Dr Jon Vernon and Prof Mark Wilcox in Leeds, but originally part of Prof SP Borriello’s collection) (17). Multilocus sequence typing (MLST) was performed directly from sequence read files using SRST2 v0.2.0 and the PubMLST *C. difficile* database (https://pubmlst.org/organisms/clostridioides-difficile/) as previously described (18, 19).

### Assembly of a new complete *C. difficile* RT 017 genome from SE Asia

To facilitate phylogenomic analysis of *C. difficile* strains from Thailand, a Thai *C. difficile* strain was selected for hybrid assembly of a closed reference genome. *C. difficile* MAR286 was a non-MDR strain as opposed to the existing MDR reference strain *C. difficile* M68 (20). The short-read sequencing was performed on an Illumina HiSeq sequencing platform (Illumina, San Diego, CA) using 150-bp paired-end chemistry to a depth of 39X coverage as previously described (19). The long-read sequencing was performed on a MinION sequencing (Nanopore, Oxford, UK). The sequencing libraries were prepared using the Ligation Sequencing Kit (SQK-LSK109) and run on a FLO-MIN106 (R9.4.1) flow cell, according to the manufacturer’s instructions, for 24 h. Hybrid assembly was performed with Unicycler v0.4.8 using a conservative mode (21). The final assembly graph was visualised and polished with Bandage v0.8.1 (22). Genome annotation was performed using the NCBI Prokaryotic Genomes Annotation Pipeline (23).

### AMR genotyping

AMR genotyping was performed as previously described (24). Briefly, all read files were interrogated against the ARGannot database (for known accessory AMR genes) with two additional genes recently described in *C. difficile, erm*(52) and *mefH* (16), and a customised *gyrA, gyrB* and *rpoB* alleles database (for known resistance-conferring point mutations) using SRST2 (18, 25). Strains that were positive for either *ermB* or *tetM* were interrogated for known transposons using SRST2 as previously described (18).

### Evolutionary analysis of *C. difficile* RT 017

To investigate the evolution and spread of *C. difficile* RT 017, core genome single nucleotide polymorphism (cgSNP) and Bayesian evolutionary analyses were performed. All PE reads were trimmed using TrimGalore v0.6.4 to remove low-quality and adapter sequences (https://github.com/FelixKrueger/TrimGalore), mapped to the genome of *C. difficile* M68 and variants identified using Snippy v4.4.5 (https://github.com/tseemann/snippy). The resulting VCF file was then screened to exclude variants occurring in the repetitive region using SnpSift v4.3t (26) and to exclude indels using VCF-annotate v0.1.15 (27). Gubbins v2.4.1 was used to identify and remove recombination sites (28). SNP-dists v0.7.0 was used to generate a pairwise cgSNP table (https://github.com/tseemann/snp-dists). Following the approach of Eyre *et al* (29) and Didelot *et al* (30), a threshold of 0-2 cgSNPs was used to determine if groups of 2 or more strains were clonally related.

To facilitate the Bayesian analysis, clonal strains were removed from the dataset leaving only one representative for each clonal cluster. Bayesian evolutionary analysis was performed using BactDating v1.0.1 (31). BactDating was run using a Gubbins recombination-adjusted phylogenetic tree from the previous analysis (1455 sites) as an input with the following settings: Markov chain Monte Carlo (MCMC) chains of 5 × 10^8^ iterations sampled every 5 × 10^5^ iterations with a 50% burn-in and a strict model with a rate of 1.4 mutations per genome per year as published by Didelot *et al* (30). Traces were inspected to ensure convergence and the effective sample sizes (ESS) for all estimated continuous variables were >200. The final Bayesian tree was annotated using iTOL v6 (32). An interactive version of the Bayesian phylogenetic tree in **Figure 2** was uploaded to Microreact and is available at https://microreact.org/project/v89tzQ8rii55PkAGF5Jo2r/64c80194 (33).

Bayesian analysis was also performed on a subset of 45 Thai *C. difficile* genomes, for which patient data and phenotypic AMR results were available (16, 34). The cgSNP analysis was performed using reference genomes listed in **Table 1** to evaluate whether the choice of reference genome had any effect on downstream analysis. A pairwise whole genome average nucleotide identity (ANI) between each *C. difficile* strain and the reference strains was generated using FastANI (35), and the results were used to compare strain relatedness with each reference.

**Table 1 –.**
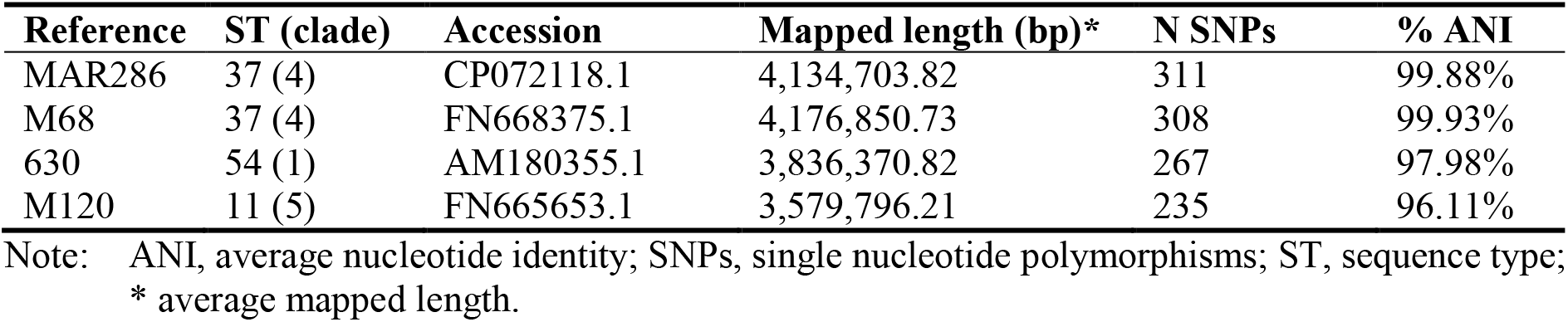
Effect of the choice of reference genome on cgSNP analysis.

### Pangenome-wide association study

The cgSNP and Bayesian analyses identified two distinct *C. difficile* RT 017 sublineages. To determine significant genetic loci associated with each lineage, all *C. difficile* genomes were assembled and a pangenome-wide association study (pan-GWAS) performed as previously described (36). Briefly, Panaroo v1.1.0 was run with default settings on the annotated *C. difficile* genomes (37), and the results used as an input for Scoary v1.6.16 to identify the significant genetic loci associated with each lineage (38).

### Assessment of virulence-associated phenotypes

We also evaluated the phenotypes associated with virulence in *C. difficile* RT 017 from the two lineages; *C. difficile* strain 1470 (ATCC 43598, non-epidemic lineage [NE]), MAR006 (epidemic lineage [E]), MAR024 (lineage E) and MAR 286 (lineage NE). First, a motility assay was performed as described by Tasteyre *et al* (39). Second, cell aggregation was assessed by measuring the optical density at 600 nm (OD_600_) of the undisturbed and disturbed 48-hour-old growth in brain heart infusion broth (40). These tests were performed with at least three biological replicates. *C. difficile* strain IS58 (RT 033, non-motile) was included as a negative control (41).

### Statistical analysis

All statistical analyses were performed using online tools by Social Science Statistics available at https://www.socscistatistics.com/. A p-value of ≤ 0.05 was considered statistically significant.

### Ethics approval

This study involved the use of de-identified patient data. It was approved by the Human Research Ethics Committee of The University of Western Australia (reference file RA/4/20/4704) and the Siriraj Institutional Review Board (protocol number 061/2558 [EC1]).

## Results

### The epidemic *C. difficile* RT 017 lineage emerged from Asia in the middle of the 20^th^ Century

To study the global population structure of *C. difficile* RT 017, cgSNP and Bayesian evolution analyses were performed on 282 non-clonal *C. difficile* RT 017 genomes collected worldwide between 1981 and 2019 (**Figure 1**). The overall median year of isolation for this dataset was 2011 (quartile range [QR]: 2008 – 2014). The median years of isolation for the three main continents were as follow; Asia, 2014 (2010 – 2016), Europe, 2010 (2006 – 2012) and North America, 2009 (2004 – 2017). Based on the Bayesian analysis (**Figure 2**), the *C. difficile* RT 017 population could be divided into two lineages: a non-epidemic lineage (NE), which could be further divided into three sublineages (NE_1_, NE_2_ and NE_3_), and an epidemic lineage (E). **Table 2** summarises 11 lineage-defining SNPs identified. Sublineages NE_1_, NE_2_ and NE_3_ consisted mainly of strains from Europe, North America and Asia, respectively, and the common ancestor of the three sublineages was estimated to have emerged in 1588 (95% confidence interval [CI]: 758 – 1858). Sublineages NE_1_ and NE_2_ split around 1860 (95% CI: 1622 – 1954). Lineage E was estimated to have split from sublineage NE_3_ around 1958 (95% CI: 1920 – 1977) and later spread globally around 1970 (95% CI: 1953 – 1983).

**Figure 1 –.**
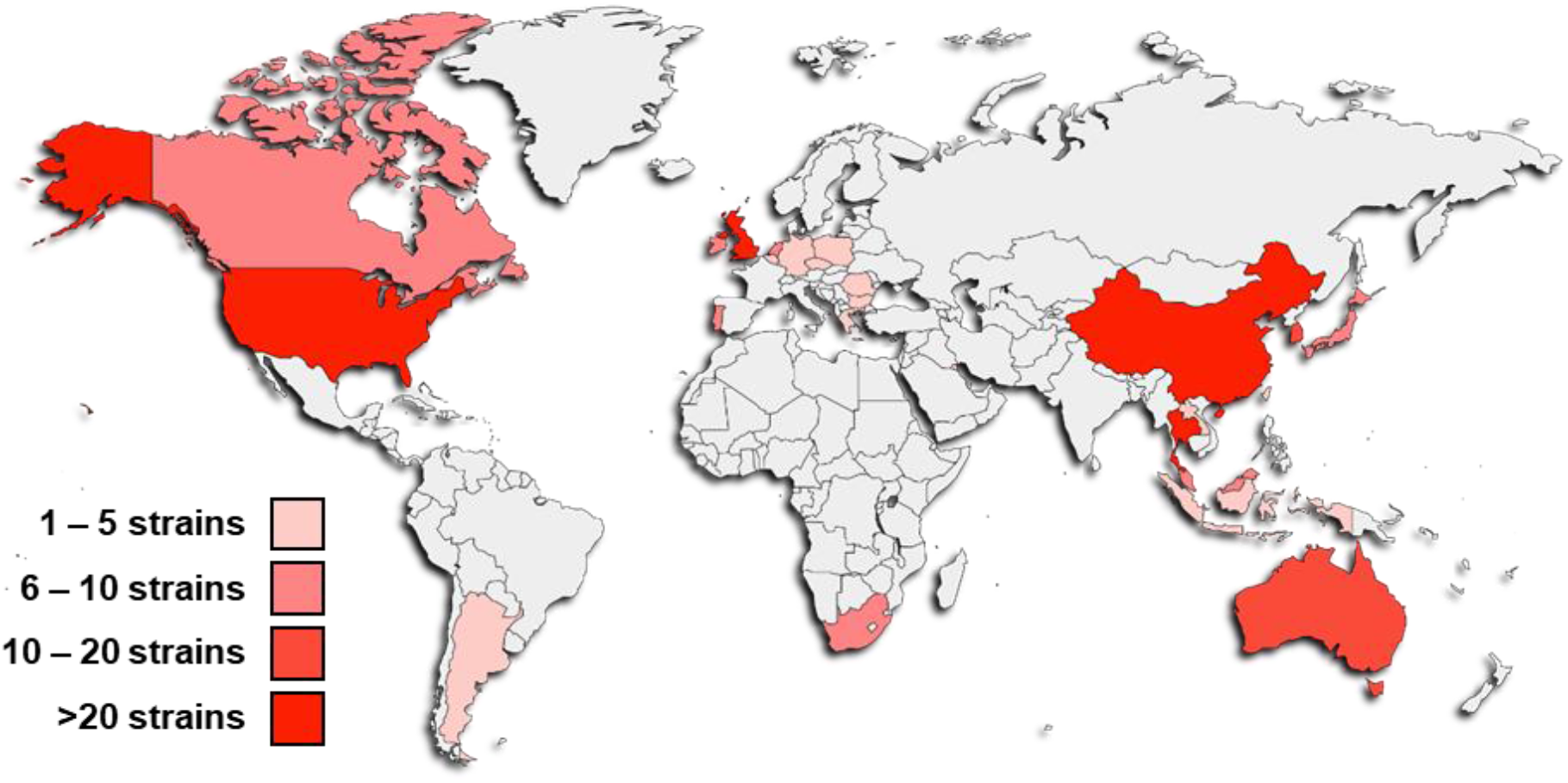
The countries of origin of the final 282 *C. difficile* RT 017 genomes. The number of genomes in each region is as follows: Africa, 6; Asia, 126; Australia, 15; Europe, 96; North America, 48; South America, 4. The world map was created by tools available at paintmaps.com.

**Figure 2 –.**
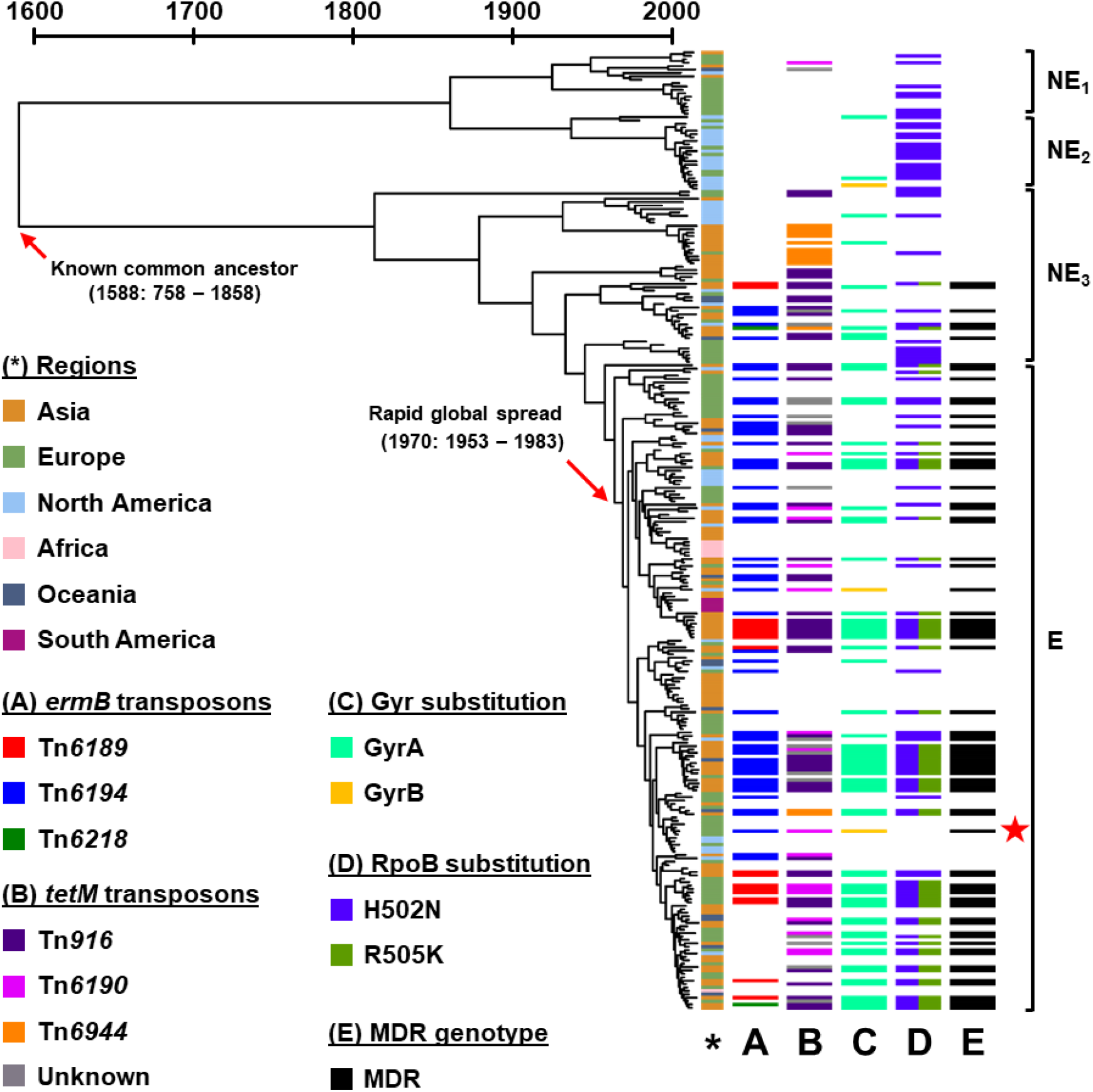
Bayesian tree of 282 non-clonal *C. difficile* RT 017 genome from around the world. *C. difficile* RT 017 population could be divided into non-epidemic (NE; sublineages NE_1_ – NE_3_) and epidemic (E) lineages. (*) refers to the region of origin for each strain. Important genotypic AMR determinants are displayed on the right (A – E). 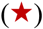 represents *C. difficile* M68, the reference genome in this analysis.

**Table 2 –.** List of lineage-defining cgSNPs.

| Position * | Strand ** | Product | $dN/dS^{***}$ | NE<sub>1</sub> | NE<sub>2</sub> | NE<sub>3</sub> | E |
| --- | --- | --- | --- | --- | --- | --- | --- |
| **Lineage NE vs lineage E** |  |  |  |  |  |  |  |
| 867,703 | F | Diguanylate kinase signalling protein | $dN$ | G<sup>†</sup> | G<sup>†</sup> | G<sup>†</sup> | T |
| **Sublineages NE<sub>1</sub> and NE<sub>2</sub> vs sublineage NE<sub>3</sub>** |  |  |  |  |  |  |  |
| 263,571 | F | FlgG | $dN$ | T<sup>†</sup> | T<sup>†</sup> | C | C |
| 480,088 | R | UvrA | $dS$ | A<sup>†</sup> | A<sup>†</sup> | G | G |
| 1,486,937 | F | Gfo/Idh/MocA family oxidoreductase | $dN$ | T<sup>†</sup> | T<sup>†</sup> | G | G |
| 1,789,300 | F | Serine O-acetyltransferase | $dS$ | C<sup>†</sup> | C<sup>†</sup> | T | T |
| 3,254,867 | R | ABC transporter | $dN$ | T<sup>†</sup> | T<sup>†</sup> | C | C |
| 3,808,791 | N/A | Non-coding region | - | G<sup>†</sup> | G<sup>†</sup> | A | A |
| **Sublineage NE<sub>1</sub> vs sublineage NE<sub>2</sub>** |  |  |  |  |  |  |  |
| 1,299,679 | F | Penicillin-binding protein 2 | $dN$ | G | T<sup>†</sup> | G | G |
| 1,486,584 | F | Gfo/Idh/MocA family oxidoreductase | $dN$ | C | T<sup>†</sup> | C | C |
| 2,928,003 | R | ABC transporter | $dN$ | G | T<sup>†</sup> | G | G |
| 3,066,957 | R | Thioether cross-link-forming SCIFF peptide maturase | $dN$ | C | T<sup>†</sup> | C | C |

\* Position on *C. difficile* M68 genome; \*\* coding strand (F, forward; R, reverse); \*\*\* nonsynonymous substitutions ( $dN$ ) and synonymous substitutions ( $dS$ ); <sup>†</sup> different from the reference genome.

### The acquisition of *ermB* was likely the driving factor of the epidemic *C. difficile* RT 017 lineage

After incorporating genotypic AMR data, an association between the acquisition of AMR genotype and the spread of *C. difficile* RT 017 was evident. Genotypically MDR *C. difficile* RT 017 strains were in the lower part of sublineage NE_3_ and lineage E, and only emerged around 1935 (95% CI: 1851 – 1969). There had been multiple acquisition events for the two most common accessory AMR determinants: *tetM* and *ermB*. The earliest acquisition of *tetM* was likely through gaining *Tn916* which occurred around 1914 (95% CI: 1799 – 1964), while the earliest acquisition of *ermB* was likely through gaining *Tn6194* which occurred around 1958 (95% CI: 1920 – 1977), notably the same timeframe as the predicted time of emergence of lineage E.

Nonsynonymous substitutions in RpoB (H502N, conferring rifamycin resistance) and in GyrA (T82I, conferring fluoroquinolone resistance) were found scattered throughout the population. In contrast, an R505K substitution in RpoB was found only in strains from sublineage NE_3_ and lineage E and was more common among Asian strains (37.2% vs 8.9%, p < 0.0001). The only European strains with an R505K substitution in RpoB were from an outbreak in Portugal (42). Three independent GyrB substitution events were identified in this dataset: two D426N substitution events in North America around 2008 (95% CI: 1998 – 2011) and 2015 (95% CI: 2012 – 2016), and one D426V substitution event in Ireland (*C. difficile* M68, the reference strain) around 2004 (95% CI: 2001 – 2005) (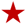 in **Figure 2**). In addition to the important AMR determinants described above, the *aac6-aph2* gene was also common among *C. difficile* RT 017, found in 73 strains in this dataset (25.9%), more commonly among Asian strains (43.4% vs 14.2%, p < 0.0001).

### The epidemic *C. difficile* RT 017 lineage expresses higher motility and lower cell aggregation

The cgSNP that differentiated between the lineages NE and E resulted in a substitution in a diguanylate kinase signalling protein, which may play role in motility and biofilm formation in *C. difficile* (40, 43). Thus, motility and cell aggregation assays were performed (**Figure 3**). Strains from lineage E had an increase in growth diameter compared to lineage NE (average diameter 7.7 vs 5.9 mm, Mann-Whitney p < 0.0001) and a slight decrease in the level of cell aggregation as shown by the lower change in OD_600_ between undisturbed and disturbed cultures (0.88 vs 0.99, Mann-Whitney p = 0.0316; for comparison, the non-motile *C. difficile* IS58 had 1.84 fold-change in OD_600_).

**Figure 3 –.**
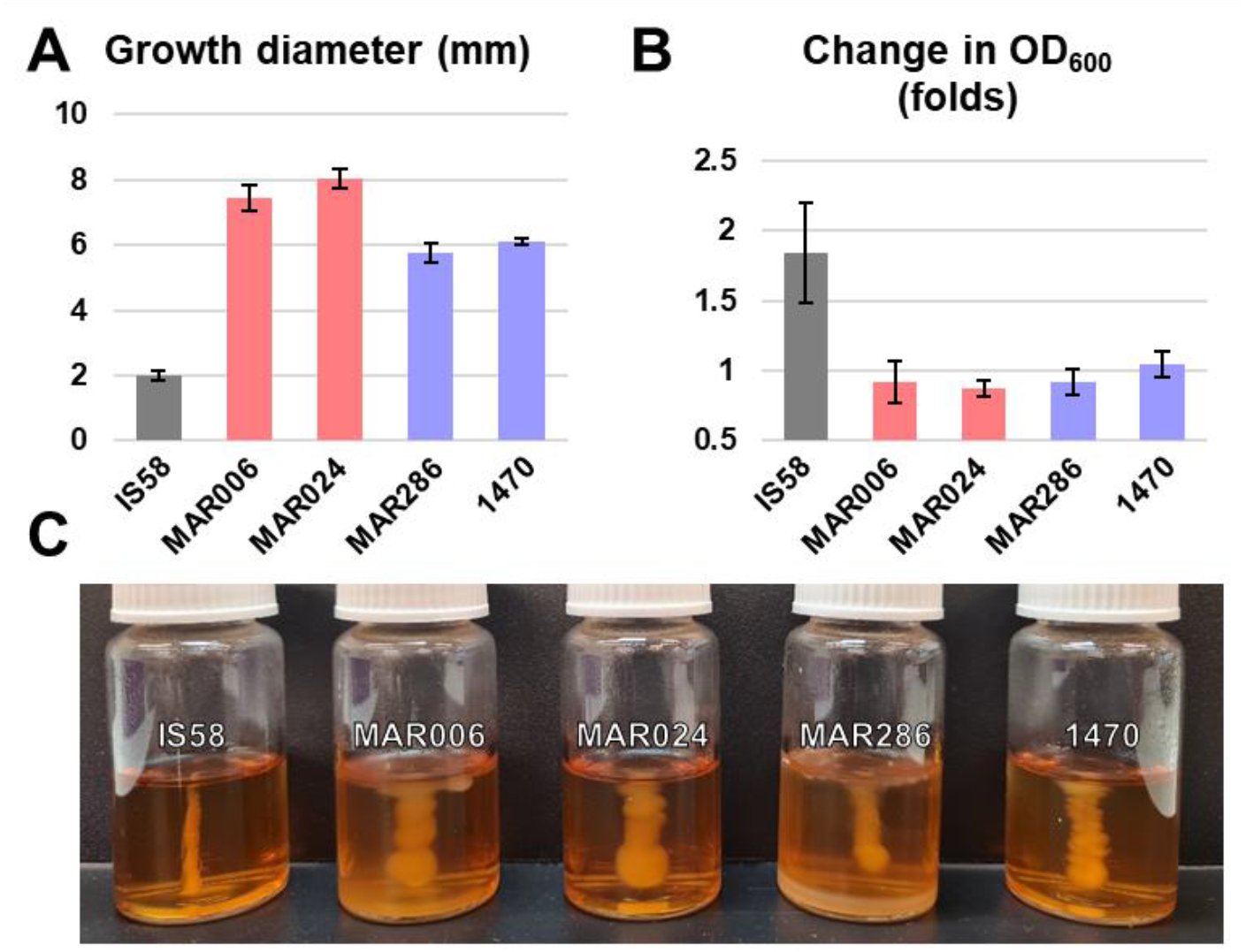
Comparison of virulence-related phenotypes between Lineages E (pink) and NE (lilac). (A) Lineage E had a larger growth diameter in semi-solid media. (B) Lineage E displayed a lower cell aggregation as measured by the difference in OD600 between undisturbed and disturbed broths. (C) The semisolid media for all tested strains. *C. difficile* IS58 (RT 033, dark grey) was used as a negative control. All error bars display 95% confidence intervals.

In addition to the lineage-specific cgSNPs (**Table 2**) and the difference in the prevalence of genotypic AMR, pan-GWAS was performed to identify other significant lineage-specific genetic loci. A total of 32,863 genes was identified in the dataset, 3,560 (10.8%) of which were found in more than 95% of strains and classified as core genes. Based on the GWAS, the locus most significantly associated with lineage E was the aminoglycoside resistance locus (containing *aac6-aph2* and a gene resembling *ant6*(Ib) [72% identity, E-value = 5.01e-157]; sensitivity 85.3%, specificity 97.8%). Apart from AMR-related loci, lineage E was associated with a truncation of the formate dehydrogenase FdhF protein (sensitivity 75.3%, specificity 97.8%). A comparison of the FdhF protein is shown in **Figure 4** (44).

**Figure 4 –.**
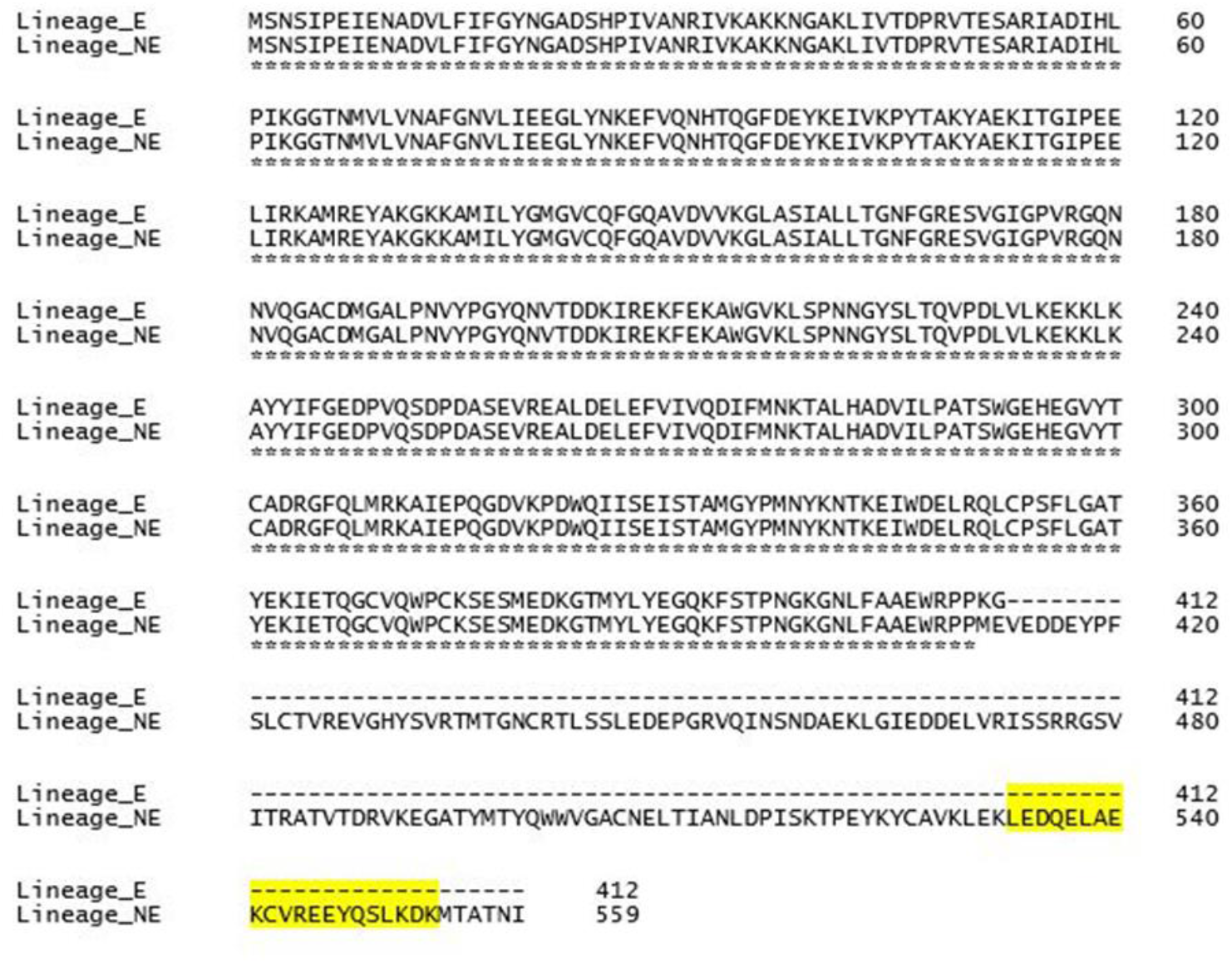
Truncation of FdhF protein in lineage E. The alignment was produced using Clustal Omega version 1.2.4 (44). The highlighted part is the predicted NAD binding site, which absent in the protein from lineage E.

### *C. difficile* RT 017 strains in Thailand were likely acquired outside of the hospital

**Table 3** compares genomes of *C. difficile* MAR286, the Thai reference genome, with *C. difficile* M68. Based on the ANI values (**Table 1**), Thai *C. difficile* strains were closest to *C. difficile* M68. Using *C. difficile* M68 as a reference resulted in the longest average mapped length, significantly longer than *C. difficile* MAR286, the second closest reference genome (p < 0.0001). Accordingly, *C. difficile* M68 was chosen as a reference for the subsequent analysis. The average number of pairwise cgSNP differences based on *C. difficile* M68 and *C. difficile* MAR286 was 0.49 SNPs (95% CI: 0.44 – 0.54). The difference between *C. difficile* strains in this study and the other two reference genomes was more pronounced resulting in a greater number of pairwise cgSNP differences compared to *C. difficile* M68: 5.42 SNPs (95% CI: 5.15 – 5.69) for *C. difficile* 630 and 9.39 SNPs (95% CI: 9.05 – 9.72) for *C. difficile* M120.

**Table 3 –.** Comparison of two *C. difficile* RT 017 reference genomes.

| Parameters | M68 | MAR286 |
| --- | --- | --- |
| Accession | FN668375.1 | CP072118.1 |
| Genome size (bp) | 4,308,325 | 4,242,261 |
| Genes | 3,983 | 3,892 |
| CDS | 3,830 | 3,761 |
| rRNA | 40 | 35 |
| tRNA | 109 | 92 |
| ncRNA | 4 | 4 |
| CRISPR array | 4 | 6 |
| % GC | 28.9% | 28.8% |
| AMR loci | *ermB* (Tn6194) [MLS<sub>B</sub>],<br>*tetM* (Tn6190) [tetracyclines],<br>D426V (GyrB) [fluoroquinolones] | *ermB* (Tn6194) [MLS<sub>B</sub>],<br>*tetM* (Tn916) [tetracyclines] |
| Pairwise ANI | 99.92% |  |

Using *C. difficile* M68 as a reference, 308 high-quality cgSNPs were identified across 45 *C. difficile* strains. The final Bayesian phylogenetic tree is shown in **Figure 5**. Based on this phylogeny, 44 *C. difficile* RT 017 strains, excluding the outlier, could be classified roughly into three groups: the oldest group (G1, n = 13), most of which were non-MDR *C. difficile* RT 017, a group of early MDR *C. difficile* RT 017 (G2, n = 15) and the most recent and rapidly expanding clade of MDR *C. difficile* RT 017 (G3, n = 16). The common ancestor of all Thai *C. difficile* RT 017 was estimated to have arisen around 1988 (95%CI: 1949 – 2000). The common ancestors of the three groups were estimated to have arisen around 1999 (1993 – 2004), 2003 (1995 – 2007) and 2012 (2009 – 2013), respectively.

**Figure 5 –.**
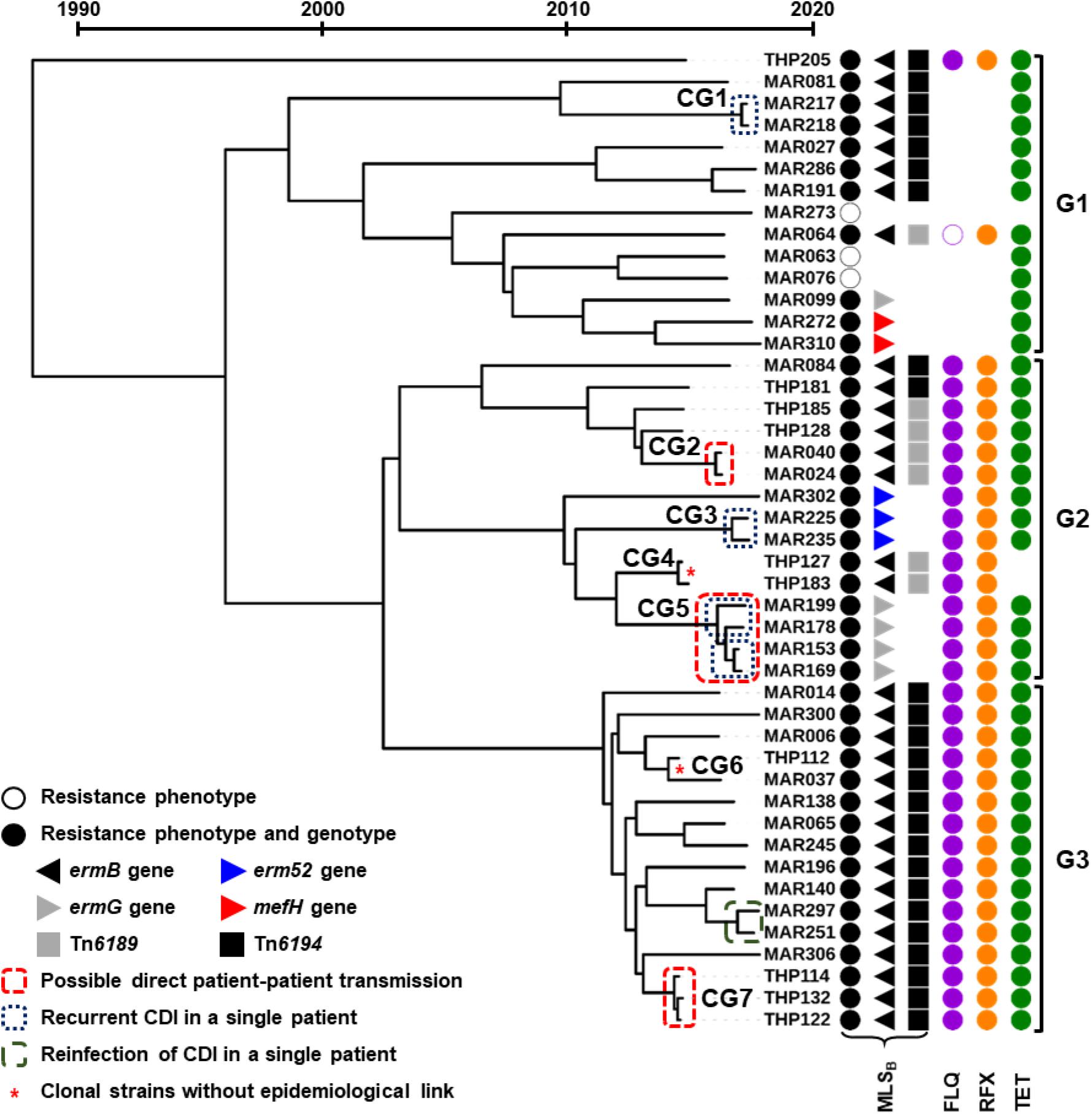
Bayesian tree of 45 Thai *C. difficile* RT 017. “THP” refers to strains isolated in 2015 and “MAR” to strains isolated in 2017 – 2018. Red boxes indicate that the patients were in the same department when the strains were isolated. Blue boxes indicated that the strains were isolated from the same patient within 2 – 8 weeks.

Seven small clonal groups (CGs) were identified across the tree (CG1 – CG7 in **Figure 5**), three of which (CG2, CG5 and CG7) were from different patients who were in the hospital during the same period, suggesting possible direct patient-patient transmission (red boxes). Two CGs (CG1 and CG3), and two small CGs in CG5, included strains that were isolated from the same patients within 2 months, suggesting recurrence CDI (blue boxes). The other two CGs (CG4 and CG6) included strains isolated from different patients without an obvious epidemiological link, one of which included strains from two specimens collected 3 years apart, suggesting contaminations in the hospital environment (red asterisks). The remaining *C. difficile* strains were non-clonal.

## Discussion

Despite being one of the most successful strains of *C. difficile*, very little is known about the evolution and spread of *C. difficile* RT 017. This study addresses this knowledge gap using high-resolution phylogenomic analyses on a comprehensive and diverse dataset of 282 global *C. difficile* RT 017 genomes. We found that the population of *C. difficile* RT 017 can be divided into two lineages, agreeing with the previous study by Cairns *et al* (14). However, data disagrees on the geographical origin of *C. difficile* RT 017. Our study suggests that *C. difficile* RT 017 may have originated in Asia, supporting the epidemiological studies (1), then spread to Europe and North America. This likely resulted from the inclusion of a few older European strains (isolated between 1981 and 1985) to reduce the gap in collection years between the two continents (p = 0.6745 in this dataset) and a large diversity of Asian strains from 11 countries and administrative regions.

Based on the difference in structure, the two lineages of *C. difficile* RT 017 were classified as non-epidemic (NE, a small number of strains with little population expansion) and epidemic (E, a larger number of strains with rapid population expansion) lineages. Although not exclusively containing strains from one continent, the NE lineage could be divided into three sublineages predominantly containing strains from Asia, Europe and North America. This suggests that the spread of *C. difficile* RT 017 between these continents had occurred since the end of the 16^th^ century. This roughly coincides with the estimated time of PaLoc acquisition ~500 years ago (45). Sublineages NE_1_ (Europe) and NE_2_ (North America) were more closely related to one another than to sublineage NE_3_ (Asia). In turn, sublineage NE_3_ was more closely related to sublineage NE_1_ than sublineage NE_2_, as demonstrated by fewer cgSNP differences (**Table 2**). Thus, the spread of *C. difficile* RT 017 likely began with population movement between Asia and Europe (1588, 95% CI: 758 – 1858) before spread from Europe to North America (1860, 95% CI: 1622 – 1954). The direction of the spread between Asia and Europe cannot be determined from this analysis, however, based on the high prevalence and diversity of clade 4 strains in Asia (9–12, 24), it is likely that *C. difficile* RT 017, as well as other strains in clade 4, originated in Asia, travelled to Europe and subsequently crossed the Atlantic to North America.

Even though *C. difficile* RT 017 could be found in at least three continents by the end of the 19^th^ century, the Bayesian analysis suggests that the epidemic lineage E emerged solely from Asia (sublineage NE_3_) following the acquisition of *ermB*-positive *Tn6194* in 1958 (95% CI: 1920 – 1977), before spreading globally in 1970 (95% CI: 1953 – 1983). The time of acquisition of the *ermB* element coincides with the introduction of clindamycin into clinical practice in the 1960s (46). This pattern of spread is similar to *C. difficile* RT 027, another epidemic strain that spread in and from North America in the early 2000s (47) driven by the acquisition of fluoroquinolone resistance in 1993/94 (47), following the widespread use of levofloxacin for community-acquired pneumonia (48). This provides supporting evidence that the use of antimicrobials and the acquisition of AMR determinants are significant in the spread of *C. difficile*. Although the prevalence of fluoroquinolone and rifamycin resistance was also high in *C. difficile*, the widespread resistance across all lineages suggests the independent acquisition of resistance after the spread of the strain.

The analyses were first performed on a small dataset of Thai clinical *C. difficile* RT 017 isolates (n = 45) with complete metadata to evaluate the performance of the pipeline. These analyses accurately identified four pairs of *C. difficile* strains isolated from the same patients, provided good correlations between AMR phenotypes and genotypes (16), as well as AMR genotypes and cgSNP population structure. When performed on the global dataset (n = 282), the analyses accurately predicted the emergence of *C. difficile* M68 (2001 – 2005), a strain from a 2003 outbreak in Ireland (20). Also, appropriate timelines for the emergence of Argentinian (1996 – 2000) and Portuguese (2003 – 2011) clusters (42, 49) were estimated, supporting the accuracy of the analyses.

Besides the aforementioned AMR genes, the epidemic lineage E was also associated with the presence of an aminoglycoside resistance locus and a truncated FdhF protein. Being a strictly anaerobic bacterium, *C. difficile* is intrinsically resistant to aminoglycosides and the presence of an additional aminoglycoside-resistance locus is unlikely to have provided any advantage to the bacterium (50). However, it may suggest that the epidemic strains were from an area with a high prevalence of aminoglycoside-resistant enteric bacteria, especially enterococci (51). Formate dehydrogenase is an enzyme involved in the reoxidation of nicotinamide adenine dinucleotide (NAD) (52). Based on the prediction by the UniProt database (53), the truncated region is the coiled-coil domain that likely serves as a binding site for NAD. Thus the truncated protein is likely non-functional, however, *C. difficile* has several pathways for oxidising NAD and the truncated FdhF may not ultimately have any effect on growth nor virulence (52). Another significant genetic variant associated with lineage E was a point substitution (W366L) on the diguanylate kinase signalling protein (**Table 2**). This protein is involved in the regulation of cyclic dimeric guanosine monophosphate (c-di-GMP) which plays a role in motility and biofilm formation (40, 43). In our preliminary assessment, strains from lineage E had increased motility and a decreased level of cell aggregation *in vitro*. Further *in vivo* studies are needed to determine how this change affects the virulence and transmissibility of the epidemic strains.

Analyses of the Thai clinical *C. difficile* strains provided information on disease transmission in the country that differs from a previous report from the UK (54). The UK study reported a cluster of closely related *C. difficile* RT 017 strains in a single hospital in London that was different to strains from other parts of the city, suggesting an intra-hospital outbreak (54). In the current study, all Thai strains were isolated in a single tertiary hospital over 4 years (2015 – 2018), however, most of them were not closely related. Overall, these strains were more related to *C. difficile* M68, a strain isolated in Ireland in a different decade (20), than to a non-epidemic strain from the same hospital. This suggests that the high prevalence of *C. difficile* RT 017 in the hospital was not due to an ongoing outbreak. Indeed, evidence of direct patient-patient transmission could be identified in only a few cases. The remaining cases acquired *C. difficile* RT 017 elsewhere, most likely from the community (55, 56).

This study also demonstrates the effect of reference genome selection on the downstream analysis (**Table 1**). The results were comparable when a reference from the same ST was used (an average of 0.49 SNPs difference, clonality cut-off point of 2 SNPs) (30). Differences became more pronounced as the reference strain became less related, suggesting that a reference genome from the same ST should be used to ensure accurate cgSNP results. With the introduction of ONT, it is now possible to assemble a complete genome of a local reference strain to maximise the accuracy of cgSNP analysis using a combination of short and long-read sequences.

A limitation of this study remained the relatively low number of early *C. difficile* RT 017 strains in general and the lack of older strains from Asia. This likely led to some uncertainty in the estimations, as reflected by wide 95% CIs, especially around the root of the Bayesian tree. Although it may be difficult to acquire old clinical strains, it may be possible to get historical strains from other sources. Soil is one promising source for ancient *C. difficile*, as it is a reservoir for *C. difficile* spores and several methods have been developed to measure the age of the soil (57), which can be used as a substitution for the collection date in a Bayesian evolutionary analysis.

In conclusion, *C. difficile* RT 017 had been circulating between Asia and Europe for centuries before spreading to North America. The epidemic lineage of *C. difficile* RT 017 emerged from Asia in the middle of the 20^th^ century following the acquisition of *ermB*. A focused investigation of contemporary *C. difficile* RT 017 in Thailand revealed that the population was highly diverse and community reservoirs/sources may have played an important role in the transmission of disease in this country.

## Acknowledgements

This work was supported, in part, by funding from The Raine Medical Research Foundation (RPG002-19) and a Fellowship from the National Health and Medical Research Council (APP1138257) awarded to D.R.K. and Edith Cowan University School of Medical and Health Sciences Research Grant Scheme awarded to D.A.C. K.I. is a recipient of the Mahidol Scholarship from Mahidol University, Thailand. This research used the facilities and services of the Pawsey Supercomputing Centre [Perth, Western Australia].

## Additional information

The **Supplementary Document** is available at DOI: 10.6084/m9.figshare.14544792.

